# Susceptibility of novel Italian rice varieties to panicle blast under field conditions

**DOI:** 10.1101/2020.04.23.057554

**Authors:** Gabriele Mongiano, Patrizia Titone, Simone Bregaglio, Luigi Tamborini

## Abstract

A panel of 48 of Italian rice varieties recently included in the Common Catalogue of Agricultural Species was evaluated for susceptibility to rice blast (*Magnaporthe oryzae* B. Couch) in open field conditions. Trials were performed under highly favourable conditions for the pathogen, and visual assessments focused on the severity and incidence of panicle blast symptoms. Only 8% of newly released varieties were classified as resistant, whereas 40% were highly susceptible. Our results confirmed that a fungicide treatment with tricyclazole reduces the disease incidence and severity and that the effect is measurable up to six weeks after treatment. A double application of tricyclazole at stem elongation and booting stage was more effective than the single application at booting. This study provides ready-to-use information to support rice growers in variety choice and planning of plant protection strategies, as well as public institutions in the emanation of guidelines for integrated pest management.

## Introduction

Over the last 25 years, about 20 new rice varieties were included annually in the Common Catalogue of Agricultural Species, half of which came from the National Register of Italy. The inclusion of a new variety in the National Register depends by its distinguishability from the other varieties of common knowledge and on its value for cultivation and use (VCU, Ministero delle Politiche Agricole Alimentari e Forestali 2014). In essence, an admissible VCU may be an improvement, when compared to the varieties of common knowledge, regarding its productivity, quality of the product, resistance to pests and diseases, as well as other technological and commercial characteristics. One of the main determinants of the spread of cultivation of a new rice variety is its resistance to major diseases. Nevertheless, susceptible varieties with valuable qualitative traits are still widely cultivated like Carnaroli and Vialone Nano, two blast-susceptible accessions traditionally used for the preparation of risotto. The breeding of disease tolerant varieties with superior technological and commercial characteristics requires great economical efforts and extended periods. It is, therefore, crucial to deepen the knowledge about the susceptibility to blast disease of current rice varieties, in order to support farmers in variety choice and related plant protection strategies.

In Italy, the most destructive rice disease is blast, caused by *Magnaporthe oryzae* B. C. Couch (anamorph *Pyricularia oryzae* Cavara) (Couch and Kohn 2002; Na et al. 2019), which determines substantial yield losses and declines in milling rice yield (Bregaglio et al. 2017; Webster and Gunnell 1992). Manifestations of the disease vary along with rice phenology. Symptoms occur primarily on the leaves (leaf blast, before flowering) and the panicle (panicle blast, PB, during ripening), the latter leading to the most considerable yield losses (Webster and Gunnell 1992), with low correlation between the two symptomatology (Biloni and Lorenzi 2002). To date, genetic resistance is the most effective control method for PB, as more than 100 genes and about 500 QTLs capable of conferring pathogen resistance have been isolated globally (Li et al. 2007; Miah et al. 2012). However, genetic resistance is rapidly overcome by the fungus within 3 to 5 years due to the high mutation rate of the pathogen populations (Oliveira-Garcia and Valent 2015; Rama Devi et al. 2015).

Five years ago, we evaluated 105 Italian varieties for PB susceptibility, concluding that, although cultivated on the 80% of the whole Italian rice area, 68% of the tested varieties were susceptible (Titone et al. 2015). Modern varieties were more resistant than old accessions, proving the constant breeding efforts towards the identification of new sources of genetic resistance. That study reported that a single fungicide application with tricyclazole during flowering was able to reduce the PB incidence by an average of 55%. However, this active substance is not anymore authorized in Europe (Reg. EC No. 1107/2009), although a new examination is underway by the competent authority. The present work aimed at evaluating the degree of susceptibility to PB in 48 new rice varieties, recently registered in the Common Catalogue and currently grown in Italy. The new rice varieties were compared with 14 reference varieties with different degrees of blast resistance (Titone et al. 2015) in dedicated field trials. Considering the possible reintroduction of tricyclazole among the admitted active substances in rice protection, single and double applications were tested, and their efficacy was also evaluated outside the period of protection (6 weeks after treatment). The information released here is ready-to-use for rice growers, to optimize plant protection strategies according to the susceptibility of the cultivated variety, as well as for public entities, which can rely on our results to plan legislative measures for integrated pest management.

## Methods

### Experimental design

Field trials were conducted during the rice-growing seasons 2015, 2016, and 2018 in “Lomellina” area (45°17’21.3“N 8°51’59.4”E, 116 meters a.s.l., Lombardy region) in the core of the Italian rice cultivation area. The site presents favourable conditions for the development of rice blast, i.e. endemic presence of the pathogen and sandy soil. Sowing of seed occurred on May 11^th^ in 2015, on May 17^th^ in 2016, and on May 18th in 2018 using a pneumatic seed drill. The experimental unit consisted of a two-meters row interspersed with spreader rows planted with the highly blast-susceptible cultivar “Deneb”. After the 3rd leaf stage the field trial was flooded until the beginning of ripening stage. A total N amount of 210 kg ha^−1^ was distributed in three applications (after 3rd leaf unfolding, during tillering, and at panicle differentiation stage), to further contribute to the development of disease, as high N rates are known to set a conducive environment for the blast pathogen (Ballini et al. 2013).

We used a strip split-plot randomized complete block design with four replications: tricyclazole was tested at the recommended rate (equivalent to 450 a.i. g ha^−1^) in single application at booting stage and in double application at stem elongation and booting; untreated plots were used as control.

The 48 rice varieties tested were registered in the period 2015-2018, with an additional group of 14 reference varieties which were known to have different blast susceptibility (Titone et al. 2015). Each variety has been tested in two out of the three years, while the reference varieties were grown in the three years. Supplementary Table S1 present the list of varieties included in the study, along with information on the shape of the caryopsis, the European market classification and the cultivated area in 2018.

### Environmental conditions

The temperatures occurred in the three years of the experiment are shown in Supplementary Figure S1. The 2015 growing season was characterized by a remarkably hot July, which caused a contraction in the duration of the vegetative cycles. The months of June and July were characterized by high temperatures (up to 37 °C) and relative humidity until the last week of July. During the middle of August, frequent rainfalls led to lower temperatures and favourable conditions for the development of the pathogen. In May and June 2016, air temperature was below-average, postponing the time of anthesis. In July and August temperatures were in line with the average, favouring a gradual recovery of the initial vegetative delay. The month of September was the hottest of the last ten years, with an average increase in maximum temperatures of about 2°C. Rainfall over the whole season was scarce, particularly in April, August and September. On average, rainfall was about 20% lower than the last decade. The 2018 growing season was warmer than the average of the last 11 years, with heavy rainfall towards the end of the season. Monthly average temperatures were always higher than the average of the last 11 years, especially in July and August, when minimum and maximum temperatures were 1 °C and 2.5 °C higher, respectively.

### Assessment of panicle blast severity and incidence

Plots were evaluated twice, after the treatment at booting stage: three and six weeks after treatment (WAT), when physiological maturity was almost reached. PB severity was assessed following the Standard Evaluation System proposed by the International Rice Research Institute (IRRI 2002). This method consists of a six scores ordinal scale from 0 to 9 based on the occurrence and extension of lesions on panicle internode and branches, also considering the number of filled grains, as follows: 0 - no visible lesions or minor lesions, 1 - lesions on several pedicels or secondary branches, 3 - lesions on a few primary branches or panicle primary axis, 5 - lesion partially around the panicle node/internode or the lower part of panicle primary axis, 7 - lesion completely around panicle node/internode or panicle primary with more than 30% of filled grains, 9 - lesions completely around panicle node/internode or primary axis with less than 30% of filled grains. We assigned a single score to the plot according to the most frequent observed symptom. Disease incidence was estimated visually as the ratio of the panicles showing PB disease over the total number of panicles in the plot (%). PB incidence at 3 and 6 WAT was used to calculate the Area Under Disease Progress Stairs (Simko and Piepho 2012), a method suited to combine multiple observations in a single value. AUDPS was used to classify varieties as Resistant (R, AUDPS ≤ 5), Moderately Resistant (MR, 5 > AUDPS ≤ 10), Moderately Susceptible (MS, 10 > AUDPS ≤ 15), Susceptible (S, AUDPS > 15). This classification has been developed consistently to previously published data on the reference varieties (Titone et al. 2015).

### Data analysis

A chi-square test of independence was performed to evaluate the relation between fungicide treatment and disease severity. The AUDPS values were analysed by Generalised Least Square model considering variety, treatment, and year effects, and accounting for heteroskedasticity due to treatment and year, by specifying their variance structures as different variances per stratum (Pinheiro and Bates 2006). The significance of the considered effects was tested using a Wald *χ*^2^ test (Fox 1997). The differences between levels of the significant effects were investigated by computing the estimated marginal means (EMM), a method to estimate the least-square means with unbalanced data, and then contrasts for the pairwise comparisons were generated (Searle et al. 1980). Estimates of the effectiveness of the treatment in reducing PB incidence and severity were calculated as the reduction in PB incidence between untreated and treated plots. Results were reported as mean reduction and relative standard error using the notation 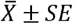. Boxplot were drawn to show the median (horizontal line), interquartile range (IQR as upper and lower hinges, i.e. 75th and 25th percentiles, respectively), and 95% confidence intervals around the median (the whiskers, calculated as following:

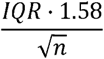

Data analysis was performed using the R statistical software (R Core Team 2017) and *nlme* R package (Pinheiro et al. 2019). Graphics were created using the ‘*ggplot’* R package (Wickham 2016).

## Results

### Disease severity

The frequency of the PB severity assessed on the sixty-two rice varieties (48 newly registered cultivars and 14 reference varieties) in the three years of the experiment is presented in Figure 1, considering the untreated control and single and double fungicide applications.

**Fig. 1.**
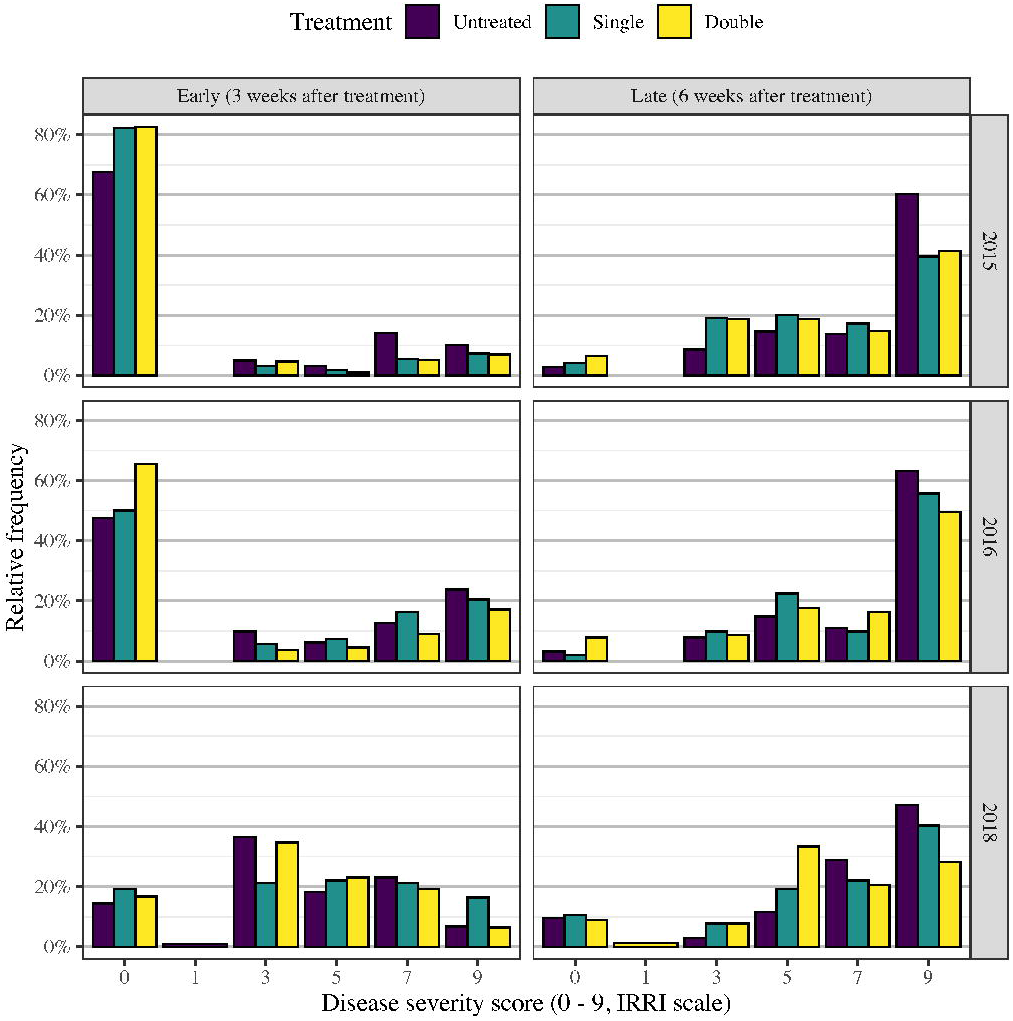
Frequency histogram of varieties by disease severity scores (IRRI scale) assigned in the field trials during the three years of experiment, divided in early and late assessments for untreated control and fungicide treatment in single and double application

The increase of PB severity from 3 to 6 WAT was clear in the three years of the experiment (Figure 1). In the early assessment, the most frequent score was 1 (lesions on several pedicels or secondary branches) in 2015 (77%) and 2016 (54%), while in 2018 it was 3 (30%, lesions on a few primary branches or panicle primary axis). In late assessments, the PB severity reached the maximum score of 9 (lesions completely around panicle node/internode or primary axis with less than 30% of filled grains), in more than 40% of the varieties in the three years. The PB symptoms onset was late in 2015 and 2016, whereas their progression was faster than in 2018, with large differences between early and late assessment. In 2018 the PB onset was earlier, with varieties evenly assigned to scores from 1 to 7 at 3 WAT; PB then slowly progressed during ripening, with little or no change for some cultivars at 6 WAT. We observed a general reduction in PB severity in the treated plots compared to the control at 3 and 6 WAT. At 3 WAT, the average ratio of varieties in which the assigned PB severity score was 0 increased from 43% in untreated plots to 51% in plots with a single application and to 55% in plots with double application of tricyclazole. In the late assessment (6 WAT), the average ratio of varieties assigned to PB severity score of 9 decreased from 57% (untreated) to 45% and 40% in plots with a single and double fungicide application, respectively. In general, fungicide treatment led to a shift in the frequencies of treated plots with respect to control, with a steep increase in the frequencies of lower PB scores. A χ^2^ test of independence performed to compare the frequencies of varieties assigned to disease severity scores among treatments indicated a significant effect, both at 3 WAT −χ^2^(10)=38.32, *p* < 0.001 – and 6 WAT −χ^2^(10) = 44.33, *p* < 0.001.

### Disease incidence

The assessment of PB incidence expressed as the ratio of infected panicles over the total number of panicles in the plot led to results similar to PB severity (Figure 2). The correlation between PB severity and incidence was evaluated by means of Spearman’s rank correlation coefficient, which highlighted a high degree of correlation both in early (*ρ* =. 85, *p* <. 05) and late (*ρ* =. 79, *p* <. 05) assessments.

**Fig. 2.**
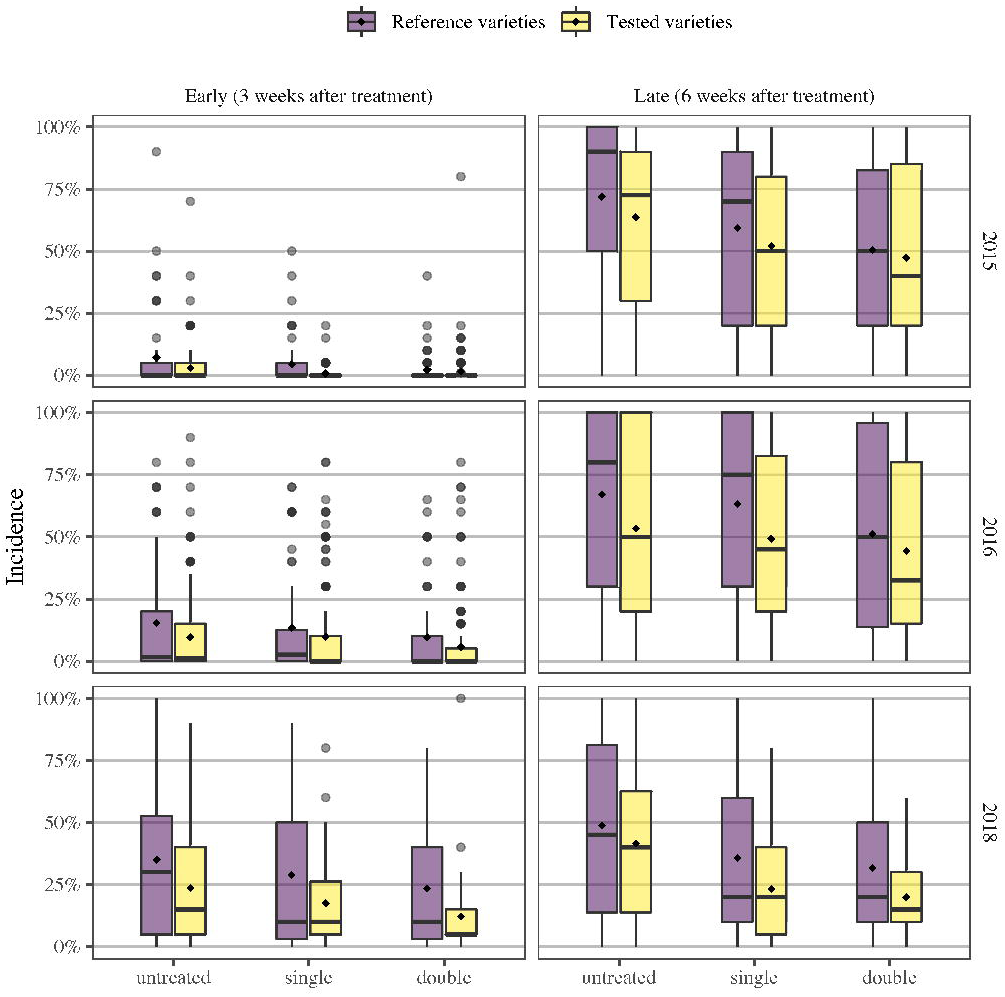
Distributions of incidence values recorded in the experiment, comparing early and late evaluation and divided into the three years of testing and the three evaluated treatments: untreated and tricyclazole in single and double application. Boxplot were drawn showing: the median (horizontal line), IQR (upper and lower hinges), and 95% confidence intervals around the median (the whiskers); the square marks indicate the group mean

As reported for PB severity, the onset of PB in 2018 was earlier than in the other two years, with high variability already at 3 WAT, when PB was generally very low in 2015 and 2016 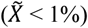. The interannual variability of PB severity and incidence can be attributed to the environmental conditions occurring during the years rice-growing season, rather than to the sample composition since comparable trends in reference and tested cultivars were outlined throughout the experiment. Another element supporting this hypothesis is that leaf blast symptoms in 2018 were much more visible in pre-flowering, especially on the extremely susceptible varieties (data not shown). The progress of the disease evaluated in untreated plots from 3 to 6 WAT was also different in the three years. Despite the low PB incidence at 3 WAT 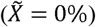, median PB incidence at 6 WAT was 80% in 2015. A slightly smoothed increasing trend was observed in 2016 (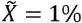 at 3 WAT, 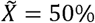 at 6 WAT), while in 2018 the progress of PB was slower, leading to a lower median incidence at 6 WAT (40%). On average, PB incidence in untreated plots increased by 82% during the three weeks between the early and late assessments in the three years. PB incidence in treated plots was generally lower than in control both in early (−61% ± 5% incidence with the double application, −25% ± 9% with single application) and late assessments (−22% ± 4% with the double application, −13% ± 4% with single application).

### Disease dynamics (AUDPS)

The AUDPS calculation allowed combining in a single value early and late assessments of PB incidence, and highlighted a comparable trend between untreated and treated plots in the three years (Figure 3). The GLS model applied to evaluate the effects of cultivar, treatment, and year on AUDPS obtained a pseudo R^2^ value of 48.5%, indicating adequate fit and large amount of explained variance. Model diagnostics, i.e. Quantile-Quantile plot and plot of Fitted values versus Standardised Residuals, are reported in Supplementary Figure S2 and S3. The significance of the considered sources of variation was then tested with Wald’s χ^2^ test (Table 1).

**Table 1.** Results of the χ^2^ test performed on AUDPS data considering the treatment (three levels, i.e. untreated, single, and double application of tricyclazole), year, and cultivar effect.

| | D.F. | $\chi^2$ | Significance |
| --- | --- | --- | --- |
| Cultivar | 62 | 2838 | < 0.001 *** |
| Treatment | 3 | 163 | < 0.001 *** |
| Year | 2 | 9.065 | 0.01075 * |
| Residuals | 1608 |  |  |

**Fig. 3.**
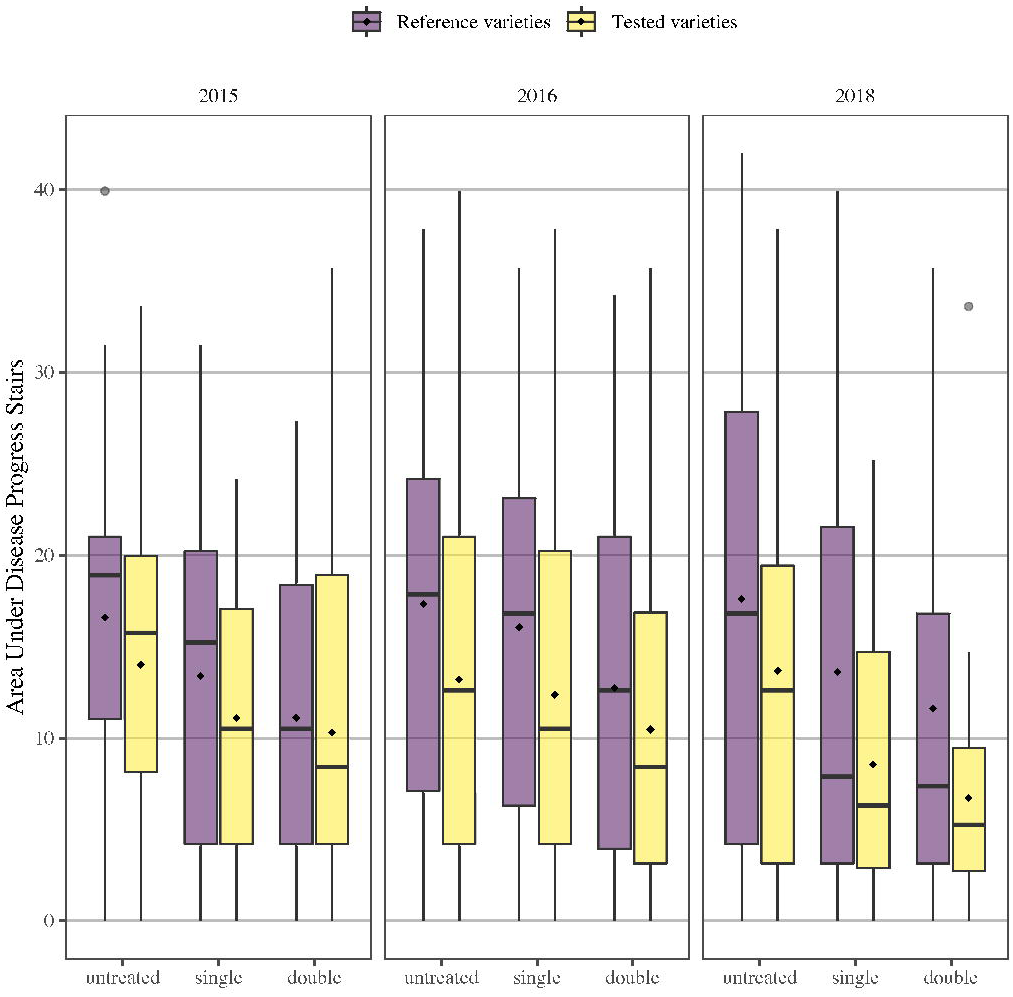
Distribution of Area Under Disease Progress Stairs (AUDPS) values over the three years of trials, comparing untreated plots with single and double application of tricyclazole, dividing reference varieties (n = 14) from tested varieties (n = 48). Boxplot were drawn showing: the median (horizontal line), IQR (upper and lower hinges), and 95% confidence intervals around the median (the whiskers); the square marks indicate the group mean

All the factors were significant (*p* < 0.05) in affecting AUDPS. We classified the 48 new varieties as Resistant (R, 0 - 5), Moderately Resistant (MR, 5.1 - 10), Moderately Susceptible (MS, 10.1 - 15), Susceptible (S, AUDPS > 15) to PB according to the average value of AUDPS in untreated plots, in agreement with published data on reference cultivars (Titone et al. 2015). We updated the former classification, which was based on a single early assessment of PB incidence, in order to consider AUDPS-based rating. Table 2 reports the estimated marginal mean, standard error, confidence intervals (95% confidence), mean Incidence (both in early and late assessments), and AUDPS calculated on untreated plots, for each variety tested. The mean AUDPS and 95% confidence intervals around the mean calculated for each cultivar in untreated plots are shown in Figure 4.

**Table 2.** AUDPS estimated marginal means (EMM) for all the levels of the variety factor. Standard error (SE), lower confidence limit (LCL), upper confidence limit (UCL), early and late PB incidence assessed in untreated plots (Inc. Early, Inc. Late), mean area under disease progress stairs calculated in untreated plots (AUDPS, untreated), and assigned tolerance class are also reported. The horizontal triple line separates the reference cultivars included in Titone et al. 2015 from those used exclusively in this study. Titone, P., Mongiano, G., & Tamborini, L. (2015). Resistance to neck blast caused by Pyricularia oryzae in Italian rice cultivars. European Journal of Plant Pathology, 142(1), 49–59. doi:10.1007/s10658-014-0588-1

| Cultivar | EMM | SE | LCL | UCL | Inc.<br>Early | Inc.<br>Late | AUDPS<br>untreated | Tolerance<br>class |
| --- | --- | --- | --- | --- | --- | --- | --- | --- |
| Clxl745 | 0.2 | 0.2 | 0 | 0.7 | 0% | 1% | 0.2 | R |
| Mare Cl | 3.5 | 0.6 | 2.4 | 4.7 | 2% | 20% | 4 | R |
| Ellebi | 7 | 1 | 5.1 | 8.8 | 6% | 37% | 9 | MR |
| Cl26 | 8.5 | 1 | 6.5 | 10.5 | 4% | 46% | 10 | MS |
| Roma | 10.3 | 1.1 | 8.2 | 12.5 | 9% | 56% | 14 | MS |
| Centauro | 12.6 | 1.1 | 10.5 | 14.7 | 9% | 56% | 14 | MS |
| Thaibonnet | 14.2 | 1.3 | 11.7 | 16.7 | 4% | 68% | 15 | MS |
| Ronaldo | 12.8 | 1.2 | 10.5 | 15.1 | 8% | 72% | 17 | S |
| Carnise Precoce | 16.6 | 1.4 | 13.8 | 19.4 | 23% | 65% | 18 | S |
| S. Andrea | 14.4 | 1.2 | 12.1 | 16.8 | 23% | 70% | 20 | S |
| Karnak | 19 | 2 | 15 | 23 | 22% | 85% | 22 | S |
| Gladio | 18 | 1.3 | 15.5 | 20.5 | 30% | 78% | 23 | S |
| Loto | 24.8 | 1.3 | 22.2 | 27.4 | 40% | 92% | 28 | S |
| Ambra | 25.2 | 1.5 | 22.2 | 28.2 | 47% | 96% | 30 | S |
| Deneb | 30.2 | 1.3 | 27.6 | 32.7 | 65% | 99% | 34 | S |
| Inov Cl | 2.8 | 0.7 | 1.4 | 4.2 | 1% | 9% | 2 | R |
| Nero Beppino | 6 | 1.6 | 2.8 | 9.1 | 0% | 18% | 3.8 | R |
| Cassiopea | 3.4 | 0.8 | 1.8 | 5 | 0% | 18% | 3.9 | R |
| Tuna | 5.6 | 1 | 3.6 | 7.7 | 2% | 21% | 4.7 | R |
| Ribaldo | 6.3 | 1.3 | 3.8 | 8.7 | 1% | 27% | 5.9 | MR |
| Gigante Vercelli | 5.9 | 1.2 | 3.6 | 8.2 | 2% | 26% | 6 | MR |
| Re Cl | 4.4 | 0.7 | 3 | 5.8 | 7% | 22% | 6.2 | MR |
| Cl 28 | 4.1 | 0.8 | 2.5 | 5.7 | 0% | 30% | 6.2 | MR |
| Carnaval | 3.1 | 1.2 | 0.8 | 5.5 | 0% | 30% | 6.2 | MR |
| Cl111 | 6.5 | 1.2 | 4.2 | 8.8 | 0% | 36% | 7.5 | MR |
| Cammeo | 5.3 | 0.9 | 3.6 | 7 | 8% | 34% | 8.7 | MR |
| Rg202 | 7.1 | 1 | 5.2 | 9 | 7% | 34% | 8.7 | MR |
| David Cl | 7.3 | 1.1 | 5.2 | 9.5 | 1% | 42% | 9.2 | MR |
| Caravaggio | 5.6 | 1.4 | 2.8 | 8.3 | 14% | 31% | 9.4 | MR |
| Filippo | 8.7 | 1.3 | 6.1 | 11.3 | 7% | 38% | 9.4 | MR |
| Agave | 7.2 | 1.1 | 5 | 9.3 | 1% | 44% | 9.5 | MR |
| Delfo | 10.1 | 1.9 | 6.4 | 13.8 | 0% | 48% | 10.1 | MS |
| Apache Red | 6.2 | 1.1 | 4.1 | 8.4 | 1% | 48% | 10.3 | MS |
| Violet Nori | 8 | 1.2 | 5.6 | 10.4 | 0% | 51% | 10.8 | MS |
| Fuoco | 8.1 | 0.9 | 6.3 | 9.9 | 6% | 48% | 11.3 | MS |
| Adone | 8.9 | 1 | 7 | 10.9 | 2% | 52% | 11.6 | MS |
| Casanova | 11.8 | 1.4 | 9.2 | 14.5 | 1% | 57% | 12.1 | MS |
| Il Cardinale | 11 | 1.6 | 7.8 | 14.1 | 21% | 39% | 12.8 | MS |
| Nerone Gold | 12.6 | 1.8 | 9.1 | 16 | 14% | 48% | 13 | MS |
| Aniride | 11.1 | 1.6 | 7.9 | 14.3 | 4% | 58% | 13 | MS |
| Fiamma | 13 | 1.1 | 10.8 | 15.2 | 5% | 58% | 13.3 | MS |
| Cl33 | 12.4 | 1.2 | 9.9 | 14.8 | 4% | 61% | 13.7 | MS |
| Leonardo | 14.7 | 1.3 | 12.1 | 17.3 | 1% | 68% | 14.4 | MS |
| Telemaco | 12.5 | 1.4 | 9.8 | 15.2 | 5% | 64% | 14.4 | MS |
| Cl a01 | 13.6 | 1.2 | 11.2 | 16 | 11% | 62% | 15.5 | S |
| Anteo | 12.4 | 1.3 | 9.8 | 15 | 1% | 74% | 15.6 | S |
| Gilda | 13.1 | 1.4 | 10.2 | 15.9 | 17% | 61% | 16.3 | S |
| Ilmoro | 15.1 | 1.2 | 12.7 | 17.5 | 0% | 79% | 16.6 | S |
| Orange Nori | 13.7 | 1.3 | 11.1 | 16.3 | 8% | 73% | 17 | S |
| Allegro | 13 | 1.1 | 10.8 | 15.3 | 2% | 81% | 17.5 | S |
| Sanluca | 16.6 | 1.2 | 14.3 | 18.8 | 1% | 87% | 18.5 | S |
| Dante | 18.1 | 2 | 14.2 | 22 | 21% | 68% | 18.6 | S |
| Samurai | 14.6 | 1.5 | 11.7 | 17.6 | 18% | 74% | 19.2 | S |
| Archimede | 13.3 | 1.5 | 10.3 | 16.4 | 16% | 77% | 19.4 | S |
| Macchiavelli | 15.7 | 1.5 | 12.7 | 18.7 | 11% | 86% | 20.4 | S |
| Aristotele | 19 | 2.1 | 14.9 | 23.1 | 18% | 81% | 20.9 | S |
| Egeo Cl | 16.3 | 1.3 | 13.7 | 18.9 | 10% | 91% | 21.1 | S |
| Reperso | 18.3 | 1.8 | 14.7 | 21.9 | 15% | 88% | 21.7 | S |
| Mirai | 17.5 | 1.7 | 14.3 | 20.8 | 18% | 88% | 22.2 | S |
| Felix | 21.9 | 1.3 | 19.2 | 24.5 | 16% | 96% | 23.6 | S |
| Marchese CI | 19.3 | 1.4 | 16.6 | 21.9 | 28% | 87% | 24.1 | S |
| Gelso | 24.5 | 1.7 | 21.1 | 27.9 | 36% | 85% | 25.3 | S |
| CI388 | 21.3 | 1.7 | 17.9 | 24.7 | 32% | 89% | 25.5 | S |

**Fig. 4.**
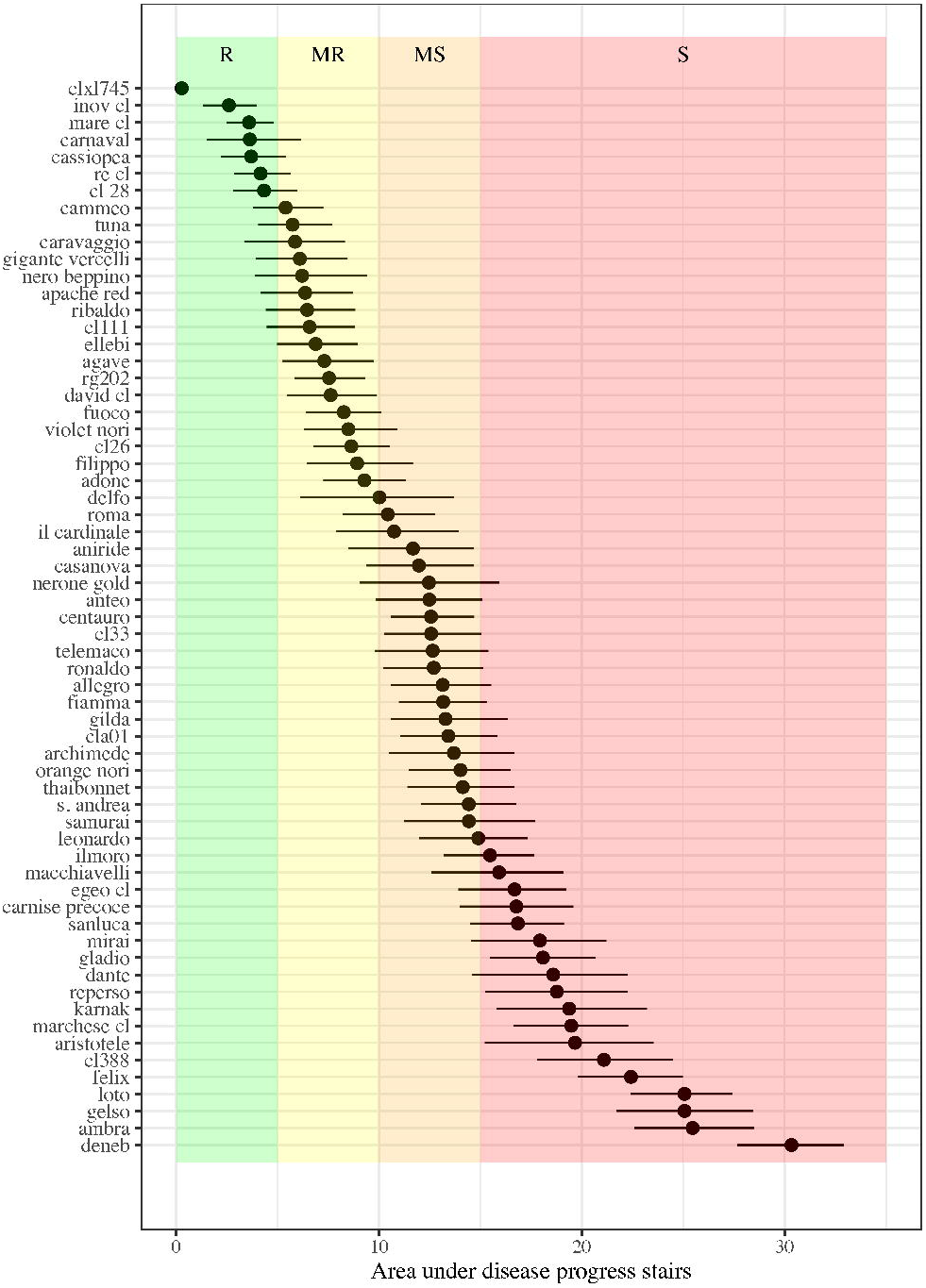
Mean values of calculated area under disease progress stairs (AUDPS) for the tested cultivars considering PB incidence assessed in untreated plots. The error bars indicate the 95% confidence interval around the mean calculated using a non-parametric bootstrap method. Plot background is color-coded to show the ranges of the four susceptibility classes used to group the cultivars as resistant (R), moderately resistant (MR), moderately susceptible (MS), and susceptible (S)

The large variability in the varietal response to PB incidence and severity was confirmed by Wald χ^2^ test that evidenced highly significant differences among cultivars (p <0.001, Table 1). While our method aimed at releasing a consistent classification with previous findings, we observed a significantly different response in some reference cultivars compared to our previous assessment (Titone et al. 2015). Roma (AUDPS 13.72) and Thaibonnet (AUDPS 14.98) showed a lower tolerance to PB compared to previous classifications, changing from MR to MS class. Karnak, Ronaldo, and S. Andrea were formerly classified as MS, while in this experiment they showed a very low degree of tolerance to PB and were classified as S, with mean AUDPS of 22.49, 16.84, and 19.6, respectively. Centauro was the only reference cultivar that showed higher tolerance to PB, passing from S to MS, with a mean AUDPS of 13.74 (early incidence 9%, late incidence 56%). The cultivar Cl 26, formerly classified as Resistant, consistently showed lower tolerance to the disease with average AUDPS of 10.54 (early Incidence 3.9%, late Incidence 46.2%). The remaining reference cultivars presented a response similar to the past and were classified consistently with published data. Cultivar Deneb, used as susceptible control and sown as “spreader row”, showed severe symptoms and a high ratio of infected panicles yet in the early assessment (average incidence 65%), and was entirely affected by the disease at 6 WAT (average incidence 99%).

During the experiment, we observed a very variable varietal response to PB (Figure 4). Only four cultivars were classified as Resistant, while 13 were MR, 13 MS, and 20 S. Among Resistant varieties, only CLXL745 (included as reference) and Inov CL showed very low incidence (<10%) both in early and late assessments. They are hybrid cultivars with late time of anthesis and long vegetative phase, which were developed in Texas and recently introduced in Italy. The other genotypes showed little to no symptoms in the early assessment but reached up to 21% PB incidence in the late assessment. MR cultivars showed an initial tolerance to disease, with PB incidence comprised between 0% and 14%, and rapid disease progress at 6 WAT, leading to PB incidence ranging from 22% to 44%. MS cultivars response in the early assessment was widely variable (0 - 21.4%), with high late PB incidence, ranging from 39.4% to 67.5%. The response of S cultivars at 3 WAT was even more variable compared to MS group (range 0.6% - 35.6%), and final PB incidence comprised between 61% and 99%.

Table 3 reports the estimated marginal means with 95% confidence intervals of the levels of treatment and year factor, with a letter-based representation of all pairwise comparisons.

**Table 3.** AUDPS estimated marginal means (EMM) for all the levels of the year and treatment effect (separated by horizontal line) with standard error (SE), and 95% lower (LCL) and upper confidence limit (UCL).

|  | EMM | SE | LCL | UCL | Subgroup |
| --- | --- | --- | --- | --- | --- |
| Year 2015 | 12.66 | 0.2291 | 12.21 | 13.11 | a |
| Year 2016 | 12.21 | 0.2264 | 11.76 | 12.65 | ab |
| Year 2018 | 11.52 | 0.3379 | 10.86 | 12.18 | b |
| Double appl. | 10.86 | 0.2577 | 10.35 | 11.36 | a |
| Single appl. | 12.00 | 0.2496 | 11.51 | 12.49 | b |
| Untreated | 13.53 | 0.2497 | 13.04 | 14.02 | c |

Single (11.27% reduction) and double (20.26% reduction) fungicide applications significantly reduced AUDPS compared to untreated plots. The differences between years were less marked, with significant differences only between 2015 (12.66) and 2018 (11.52), but not with 2016 (12.21).

## Discussion

Genetic tolerance to blast fungus *Magnaporthe oryzae* B. C. Couch has been the primary driver of Italian rice breeding throughout the twentieth century and until today (Faivre-Rampant et al. 2010; Mantegazza et al. 2008; Mongiano et al. 2018; Spada et al. 2004). PB is the most devastating symptomatology, occurring during early grain filling stage on the panicle and panicle neck node (Webster and Gunnell 1992). It causes direct yield losses by impairing the translocation of carbohydrates from the vegetative parts to grains (Koutroubas et al. 2009). Genetic resistance is undoubtedly the most effective method of control to date (Deng et al. 2017; Miah et al. 2012; Ramalingam et al. 2003; Xiao et al. 2017; Zhao et al. 2018), and recent breakthroughs in genetic improvement have opened up a plethora of new options in this regard (Wang et al. 2016; Weeks et al. 2016). Besides fungicide treatments, rice growers can choose different crop management strategies to contain the disease like, e.g. reducing nitrogen inputs and modulating the time of sowing (Ballini et al. 2013). However, the knowledge of the degree of susceptibility of the available rice varieties is the fundamental prerequisite to plan management strategies to optimize farmers actions.

In a previous study (Titone et al. 2015) we examined the tolerance of 105 Italian cultivars to rice blast disease and this characterization had been used by regional authorities to issue variety-specific guidelines for the sustainable use of plant protection products under the Water Framework Directive 2000/60/EC (European Council European Parliament 2000; Regione Piemonte 2016). Since 2015, many new Italian varieties have been introduced, and there is the need to extend the classification to the new releases. Furthermore, the most widely used active compound for PB control, tricyclazole, is currently not authorized in Europe, unless readmission following evaluation according to Reg. (EC) No. 1107/2009 could be possible.

During the three years of experiment, we observed a considerable impact of blast disease, favoured by the extremely conducive environment established in the experimental trial, i.e. high nitrogen rates applied in sandy soil, late sowing, and the presence of “spreader rows” sown with the highly susceptible cultivar Deneb. At the first assessment (3 WAT, grain filling stage), we generally observed low PB severity and incidence, while in the second assessment (6 WAT, physiological maturity) PB reached high severity and incidence. The temporal dynamics of the epidemic were similar in 2015 and 2016 and different from 2018. The main differences concerned the time of disease onset and the rate of increase of PB incidence and severity from early to late assessment. While in 2015 and 2016 PB symptoms mainly developed after the first assessment at 3 WAT, in 2018 disease onset was earlier, with also moderate infection on the leaves before flowering. Nonetheless, the PB incidence at 6 WAT in 2018 was lower compared to 2015 and 2016. This evidence disagrees with several studies, which report a higher final incidence associated with early symptoms (Filippi and Prabhu 1997; Long et al. 2001; Nasruddin and Amin 2012). However, the environmental conditions in our study were very different from those explored in the cited literature, and other findings suggest that in Italy leaf blast does not consistently correlate with PB (Biloni and Lorenzi 2002). Similar trends in disease development were reproduced in another study focusing on the Italian rice area, where a large variability of leaf and PB severity was due to inter-annual meteorological conditions (Bregaglio et al. 2016).

Compared to the experiments performed in 2011 and 2012 (Titone et al. 2015), where tricyclazole efficacy was evaluated as a single application at a rate of 450 a.i. g/ha, the substance was less effective in reducing PB, with comparable results only with a double application (average PB incidence reduction of 25% with single and 61% with the double application at 3 WAT, compared to 55% in the previous study). A possible explanation may be is that in 2015-2018 the PB incidence was much lower than in the previous study (43% mean incidence at 3 WAT) (Titone et al. 2015).

Our study confirmed that correct timing in the fungicide application is critical in reducing PB impact and must consider both the variety-specific phenological development and the development of the pathogen. The effect of a single treatment with tricyclazole was, in fact, very variable both between years and between cultivars within years. Results with the double application were more stable, having provided a more extended protection period also for very early or very late cultivars and therefore better control of the disease.

The experimental design forced us to simultaneously apply fungicides to all varieties without considering the differences in phenology and, therefore, the period of greater susceptibility to the disease (i.e. booting). This is a limitation, given that targeting the timing of fungicide application for each variety could have maximized the treatment effect, even if it would have complicated the execution of the experiment. A major improvement in the present study is the execution of a late assessment at 6 WAT, aimed at better characterizing PB incidence and to provide an indication of disease progression around maturity, outside the protection period granted by the tricyclazole treatment. Data indicated that, when favourable conditions for disease development were present, disease control had a long-lasting effect probably connected to a reduction of the inoculum source and thus, the occurrence of secondary infections. This has also been observed in *in vitro* studies (Kunova et al. 2012) and suggested by the results of the double application, which provided a more extended protection period and prevented the spread of inoculum sources.

One of the aspects that remains unexplored here is the impact of the treatment in reducing grain and milling yield losses. PB is indeed known to affect both the quantity and quality of rice yield, with a significant impact on both the gross saleable product and the selling price, as rice mills considerably reduce the purchase price of paddy rice stocks with low processing yields. Our results, however, indicate that fungicide application could be more economically justified with high-value varieties with low blast resistance, a widespread situation in the Italian rice varietal landscape (Titone et al. 2015).

Similarly to our previous study, more than 65% of the tested cultivars were found to be moderately susceptible to susceptible, and only 4 (about 8%) were classified as resistant. Therefore, even recently released varieties show low tolerance to blast disease, and this is still a critical issue for rice cultivation in Italy. Moreover, given that some reference varieties were more susceptible in our experiment than in the past, we could hypothesize an evolution of *M. oryzae* strains leading to the breakdown of some partial resistances, which is a phenomenon consistently observed in the fields (Oliveira-Garcia and Valent 2015; Rama Devi et al. 2015).

## Conclusions

The most effective, environmentally-friendly and economical strategy to control rice blast is the adoption of host resistance. However, plant breeding is an expensive and time-consuming activity requiring many years for pyramiding multiple blast resistance genes and release a new cultivar. At the same time, the pathogen population is capable of overcoming resistance genes in a relatively short period. Most of the high value, traditional Italian rice varieties, as well as the majority of recently releases confirmed to be susceptible to the disease. When genetic resistance is not an option, proper agronomic management, optionally coupled with fungicide application is, therefore, crucial to safeguard agricultural production and grower’s income in the years to come.

## Acknowledgements

Corteva (DU Pont, DOW Agrosciences) for financial support for the research activities. AgriDigit-Agromodelli project (DM n. 36502 of 20/12/2018), funded by the Italian Ministry of Agricultural, Food and Forestry Policies and Tourism.

## References

Ballini, E., Nguyen, T. T., & Morel, J.-B. (2013). Diversity and genetics of nitrogen-induced susceptibility to the blast fungus in rice and wheat. Rice, 6(1), 32. doi:10.1186/1939-8433-6-32

Biloni, M., & Lorenzi, E. (2002). Relation between leaf and neck blast resistance in Italian rice varieties [*Oryza sativa* L. - Lombardy]. Presented at the Atti delle Giornate Fitopatologiche (Italy), Pavia, Italy.

Bregaglio, S., Titone, P., Cappelli, G., Tamborini, L., Mongiano, G., & Confalonieri, R. (2016). Coupling a generic disease model to the WARM rice simulator to assess leaf and panicle blast impacts in a temperate climate. European Journal of Agronomy, 76, 107–117. doi:10.1016/j.eja.2016.02.009

Bregaglio, S., Titone, P., Hossard, L., Mongiano, G., Savoini, G., Piatti, F. M., et al. (2017). Effects of agro-pedo-meteorological conditions on dynamics of temperate rice blast epidemics and associated yield and milling losses. Field Crops Research, 212, 11–22. doi:10.1016/j.fcr.2017.06.022

Couch, B. C., & Kohn, L. M. (2002). A multilocus gene genealogy concordant with host preference indicates segregation of a new species, *Magnaporthe oryzae*, from *M. grisea*. Mycologia, 94(4), 683–693. doi:10.1080/15572536.2003.11833196

Deng, Y., Zhai, K., Xie, Z., Yang, D., Zhu, X., Liu, J., et al. (2017). Epigenetic regulation of antagonistic receptors confers rice blast resistance with yield balance. Science, 355(6328), 962–965. doi:10.1126/science.aai8898

European Council, European Parliament. (2000). Directive 2000/60/EC of 23 October 2000 establishing a framework for Community action in the field of water policy. Official Journal, L 327, 1–73. https://eur-lex.europa.eu/legal-content/EN/TXT/HTML/?uri=CELEX:32000L0060&from=EN

Faivre-Rampant, O., Bruschi, G., Abbruscato, P., Cavigiolo, S., Picco, A. M., Borgo, L., et al. (2010). Assessment of genetic diversity in Italian rice germplasm related to agronomic traits and blast resistance (*Magnaporthe oryzae*). Molecular Breeding, 27(2), 233–246. doi:10.1007/s11032-010-9426-0

Filippi, M. C., & Prabhu, A. S. (1997). Integrated Effect of Host Plant Resistance and Fungicidal Seed Treatment on Rice Blast Control in Brazil. Plant Disease, 81(4), 351–355. doi:10.1094/PDIS.1997.81.4.351

Fox, J. (1997). Applied regression analysis, linear models, and related methods. Thousand Oaks, CA: Sage Publications, Inc.

IRRI. (2002). Standard Evaluation System for Rice (SES). International Rice Research Institute. http://www.knowledgebank.irri.org/images/docs/rice-standard-evaluation-system.pdf

Koutroubas, S. D., Katsantonis, D., Ntanos, D. A., & Lupotto, E. (2009). Blast fungus inoculation reduces accumulation and remobilization of pre-anthesis assimilates to rice grains. Phytopathologia Mediterranea, 48(2), 240–252.

Kunova, A., Pizzatti, C., & Cortesi, P. (2012). Impact of tricyclazole and azoxystrobin on growth, sporulation and secondary infection of the rice blast fungus, *Magnaporthe oryzae*. Pest Management Science, 69(2), 278–284. doi:10.1002/ps.3386

Li, Y. B., Wu, C. J., Jiang, G. H., Wang, L. Q., & He, Y. Q. (2007). Dynamic analyses of rice blast resistance for the assessment of genetic and environmental effects. Plant Breeding, 126(5), 541–547. doi:10.1111/j.1439-0523.2007.01409.x

Long, D. H., Correll, J. C., Lee, F. N., disease, D. T. P., 2001. (2001). Rice blast epidemics initiated by infested rice grain on the soil surface. Am Phytopath Society, 85(6), 612–616. doi:10.1094/PDIS.2001.85.6.612

Mantegazza, R., Biloni, M., Grassi, F., Basso, B., Lu, B. R., Cai, X. X., et al. (2008). Temporal Trends of Variation in Italian Rice Germplasm over the Past Two Centuries Revealed by AFLP and SSR Markers. Crop Science, 48(5), 1832. doi:10.2135/cropsci2007.09.0532

Miah, G., Rafii, M. Y., Ismail, M. R., Puteh, A. B., Rahim, H. A., Asfaliza, R., & Latif, M. A. (2012). Blast resistance in rice: a review of conventional breeding to molecular approaches. Molecular Biology Reports, 40(3), 2369–2388. doi:10.1007/s11033-012-2318-0

Ministero delle Politiche Agricole Alimentari e Forestali. (2014). Criteri e procedure tecniche per l’iscrizione al Registro Nazionale di varietà di riso. Gazzetta Ufficiale della Repubblica Italiana - Serie Generale n. 91. http://scs.entecra.it/proveiscrizioni/criteri/riso/Criteri-Riso-GU-91-18-4-2014.pdf

Mongiano, G., Titone, P., Tamborini, L., Pilu, R., & Bregaglio, S. (2018). Evolutionary trends and phylogenetic association of key morphological traits in the Italian rice varietal landscape. Scientific Reports, 8(1), 13612. doi:10.1038/s41598-018-31909-1

Na, G., Aa, K., V Ga, M., La, R., & Sb, R. (2019). Morphological and molecular characterization of *Magnaporthe oryzae* B. Couch, inciting agent of rice blast disease. Madras Agricultural Journal, 106(Spl), 1–7. doi:10.29321/MAJ.2019.000256

Nasruddin, A., & Amin, N. (2012). Effects of Cultivar, Planting Period, and Fungicide Usage on Rice Blast Infection Levels and Crop Yield. Journal of Agricultural Science, 5(1), 160–167. doi:10.5539/jas.v5n1p160

Oliveira-Garcia, E., & Valent, B. (2015). How eukaryotic filamentous pathogens evade plant recognition. Current Opinion in Microbiology, 26, 92–101. doi:10.1016/j.mib.2015.06.012

Pinheiro, J., & Bates, D. (2006). Mixed-effects models in S and S-PLUS. Springer Science \& Business Media.

Pinheiro, J., Bates, D., DebRoy, S., Sarkar, D., R Core Team. (2019). *nlme*: Linear and Nonlinear Mixed Effects Models.

R Core Team. (2017). R: A Language and Environment for Statistical Computing. http://www.R-project.org/

Rama Devi, S. J. S., Singh, K., Umakanth, B., Vishalakshi, B., Renuka, P., Vijay Sudhakar, K., et al. (2015). Development and Identification of Novel Rice Blast Resistant Sources and Their Characterization Using Molecular Markers. Rice Science, 22(6), 300–308. doi:10.1016/j.rsci.2015.11.002

Ramalingam, J., Vera Cruz, C. M., Kukreja, K., Chittoor, J. M., Wu, J. L., Lee, S. W., et al. (2003). Candidate defense genes from rice, barley, and maize and their association with qualitative and quantitative resistance in rice. Molecular Plant-Microbe Interactions, 16(1), 14–24. doi:10.1094/MPMI.2003.16.1.14

Regione Piemonte. (2016). Deliberazione della Giunta Regionale, n. 32-2952. Piano di Gestione del Distretto idrografico del Fiume Po 2015-2021 - disposizioni attuative delle misure regionali per la riduzione dei prodotti fitosanitari nelle acque attraverso l’implementazione del Piano d‘Azione Nazionale per l’uso sostenibile dei prodotti fitosanitari. Area a vocazione risicola. http://www.regione.piemonte.it/governo/bollettino/abbonati/2016/08/attach/dgr_02952_930_22022016.pdf

Searle, S. R., Speed, F. M., & Milliken, G. A. (1980). Population marginal means in the linear model: An alternative to least squares means. The American Statistician, 34, 216–221.

Simko, I., & Piepho, H.-P. (2012). The Area Under the Disease Progress Stairs: Calculation, Advantage, and Application. Phytopathology, 102(4), 381–389. doi:10.1094/phyto-07-11-0216

Spada, A., Mantegazza, R., Biloni, M., Caporali, E., & Sala, F. (2004). Italian rice varieties: historical data, molecular markers and pedigrees to reveal their genetic relationships. Plant Breeding, 123(2), 105–111. doi:10.1046/j.1439-0523.2003.00950.x

Titone, P., Mongiano, G., & Tamborini, L. (2015). Resistance to neck blast caused by *Pyricularia oryzae* in Italian rice cultivars. European Journal of Plant Pathology, 142(1), 49–59. doi:10.1007/s10658-014-0588-1

Wang, F., Wang, C., Liu, P., Lei, C., Hao, W., Gao, Y., et al. (2016). Enhanced Rice Blast Resistance by CRISPR/Cas9-Targeted Mutagenesis of the ERF Transcription Factor Gene OsERF922. PLoS ONE, 11(4), e0154027–18. doi:10.1371/journal.pone.0154027

Webster, R. K., & Gunnell, P. S. (1992). Compendium of rice diseases. (pp. 14–16). American Phytopathological Society.

Weeks, D. P., Spalding, M. H., & Yang, B. (2016). Use of designer nucleases for targeted gene and genome editing in plants. Plant Biotechnology Journal, 14(2), 483–495. doi:10.1111/pbi.12448

Wickham, H. (2016). ggplot2: Elegant Graphics for Data Analysis. Springer-Verlag New York. http://ggplot2.tidyverse.org/

Xiao, N., Wu, Y., Pan, C., Yu, L., Chen, Y., Liu, G., et al. (2017). Improving of Rice Blast Resistances in *Japonica* by Pyramiding Major R Genes. Frontiers in plant science, 7(6258), 340–10. doi:10.3389/fpls.2016.01918

Zhao, H., Wang, X., Jia, Y., Minkenberg, B., Wheatley, M., Fan, J., et al. (2018). The rice blast resistance gene Ptr encodes an atypical protein required for broad-spectrum disease resistance. Nature communications, 1–12. doi:10.1038/s41467-018-04369-4

