## Supplementary Material for "Susceptibility of novel Italian rice varieties to panicle blast under field conditions"

Supplementary Material - European Journal of Plant Pathology

Gabriele Mongiano, Patrizia Titone, Simone Bregaglio, Luigi Tamborini

#### Contents

**1 Methods** **S2**

1.1 List of cultivars included in the study . . . S2

**2 Environmental conditions** **S4**

2.1 Temperatures . . . S4

**3 Results** **S5**

3.1 Model diagnostics . . . S5

### 1 Methods

#### 1.1 List of cultivars included in the study

Supplementary Table S1: List of cultivars included in the study with indication of year of release as inclusion in the Common Catalogue or as the date of application for Plant Variety Rights, Grain shape, Market Classification of Grain Shape, and cultivated area in 2018 (data obtained from Ente Nazionale Risi).

| Variety | Year of release | Grain shape | Market Classification | Cultivated Area (ha) |
|---|---|---|---|---|
| Adone | 2008 | half spindle-shaped | long A | 0 |
| Agave | 2010 | spindle-shaped | long A | 0 |
| Allegro | 2016 | half spindle-shaped | long A | 275 |
| Aniride | 2013 | semi-round | | |
| Anteo | — | half spindle-shaped | | |
| Apache Red | 2017 | spindle-shaped | | |
| Archimede | 2012 | half spindle-shaped | long A | 200 |
| Ariosto Cl | 2016 | long spindle-shaped | long B | 21 |
| Aristotele | 2008 | semi-round | round | 0 |
| Cammeo | 2012 | half spindle-shaped | long A | 7342 |
| Caravaggio | 2012 | half spindle-shaped | long A | 3727 |
| Carnaval | 2016 | half spindle-shaped | long A | 190 |
| Casanova | 2016 | half spindle-shaped | long A | 55 |
| Cassiopea | 2012 | long spindle-shaped | long B | 0 |
| Cl 28 | 2017 | long spindle-shaped | long B | 1453 |
| Cl111 | 2016 | long spindle-shaped | long B | 853 |
| Cl33 | 2017 | spindle-shaped | long A | 15 |
| Cl388 | 2018 | half spindle-shaped | long A | 108 |
| Cla01 | 2018 | half spindle-shaped | long A | 148 |
| Dante | 2016 | half spindle-shaped | long A | 802 |
| David Cl | 2014 | long spindle-shaped | long B | 0 |
| Delfo | 2018 | long spindle-shaped | long B | 3 |
| Egeo Cl | 2014 | semi-round | long A | 3 |
| Felix | 2017 | semi-round | round | 141 |
| Fiamma | 2017 | long spindle-shaped | long B | 76 |
| Filippo | 2012 | semi-round | round | 0 |
| Fuoco | 2017 | semi-round | | |
| Gelso | 2017 | long spindle-shaped | long B | 2 |
| Gigante Vercelli | 2016 | half spindle-shaped | long A | 8 |
| Gilda | 2017 | half spindle-shaped | medium | 0 |
| Il Cardinale | 2016 | half spindle-shaped | medium | 2 |
| Ilmoro | 2014 | half spindle-shaped | medium | 0 |
| Inov Cl | 2018 | long spindle-shaped | long B | 30 |
| Leonardo | 2017 | half spindle-shaped | long A | 1677 |
| Macchiavelli | — | semi-round | | |
| Marchese Cl | 2018 | half spindle-shaped | long A | 0 |
| Mirai | 2017 | semi-round | round | 439 |
| Nero Beppino | 2014 | half spindle-shaped | medium | 0 |
| Nerone Gold | 2018 | half spindle-shaped | medium | 86 |
| Orange Nori | 2014 | half spindle-shaped | medium | 11 |
| Re Cl | 2018 | half spindle-shaped | long A | 0 |
| Reperso | 2016 | half spindle-shaped | long A | 1 |
| Rg202 | 2017 | half spindle-shaped | long A | 6 |

Supplementary Table S1: List of cultivars included in the study with indication of year of release as inclusion in the Common Catalogue or as the date of application for Plant Variety Rights, Grain shape, Market Classification of Grain Shape, and cultivated area in 2018 (data obtained from Ente Nazionale Risi). *(continued)*

| Variety | Year of release | Grain shape | Market Classification | Cultivated Area (ha) |
|---|---|---|---|---|
| Ribaldo | 2016 | half spindle-shaped | long A | 134 |
| Samurai | 2018 | half spindle-shaped | long A | 96 |
| Sanluca | 2017 | semi-round | long A | 81 |
| Telemaco | 2017 | half spindle-shaped | long A | 1896 |
| Tuna | 2014 | half spindle-shaped | long A | 0 |
| Valente | 2018 | spindle-shaped | long A | 10 |
| Violet Nori | 2014 | half spindle-shaped | medium | 11 |

#### 2 Environmental conditions

##### 2.1 Temperatures

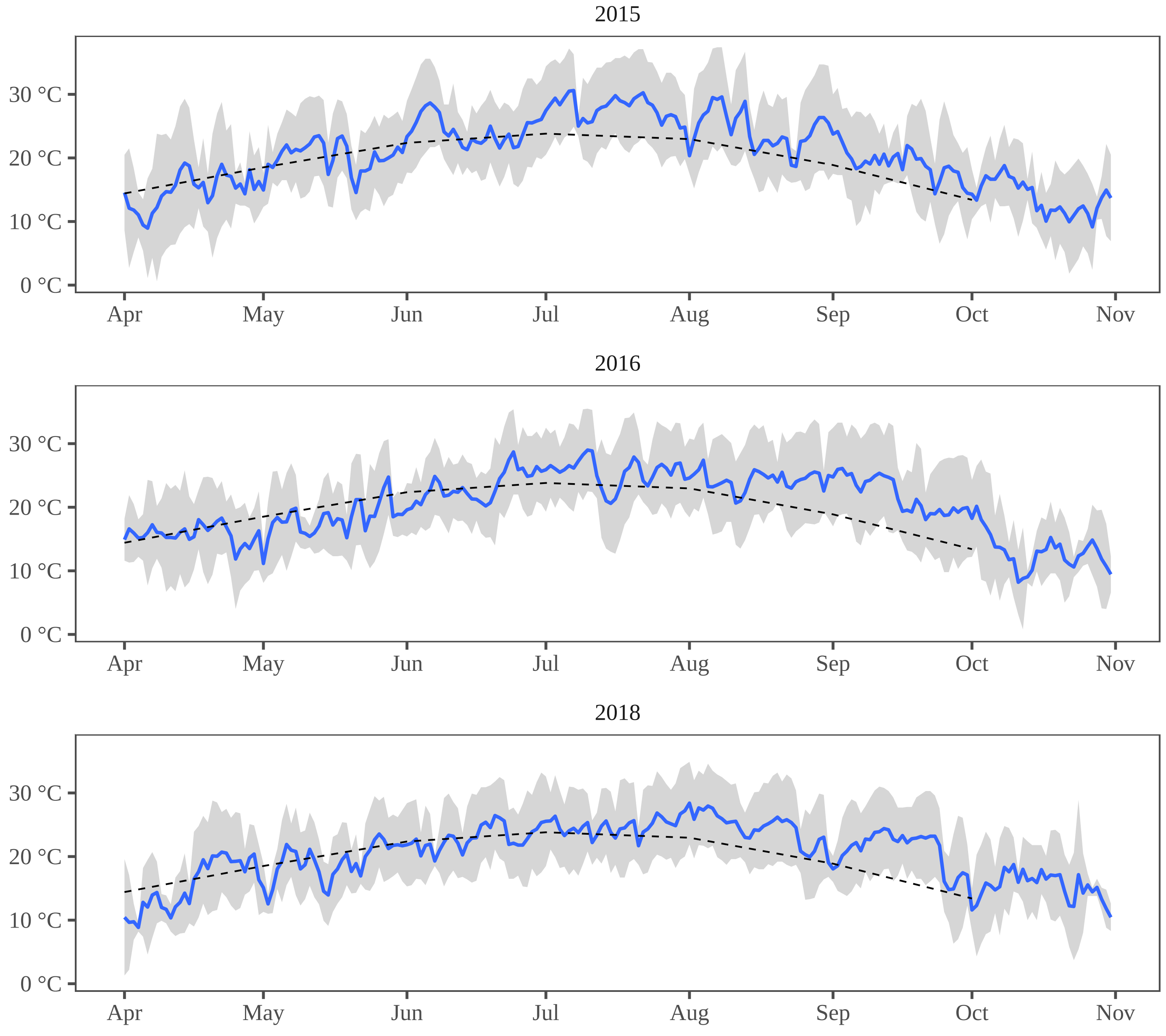


Supplementary Figure S1: Temperatures in the three years of experiment. The gray area shows the range between minimum and maximum temperatures, while the blue line is the average temperature. Dotted line represent the average temperatures of period 2007-2017.

### 3 Results

#### 3.1 Model diagnostics

##### 3.1.1 Quantile-quantile plot

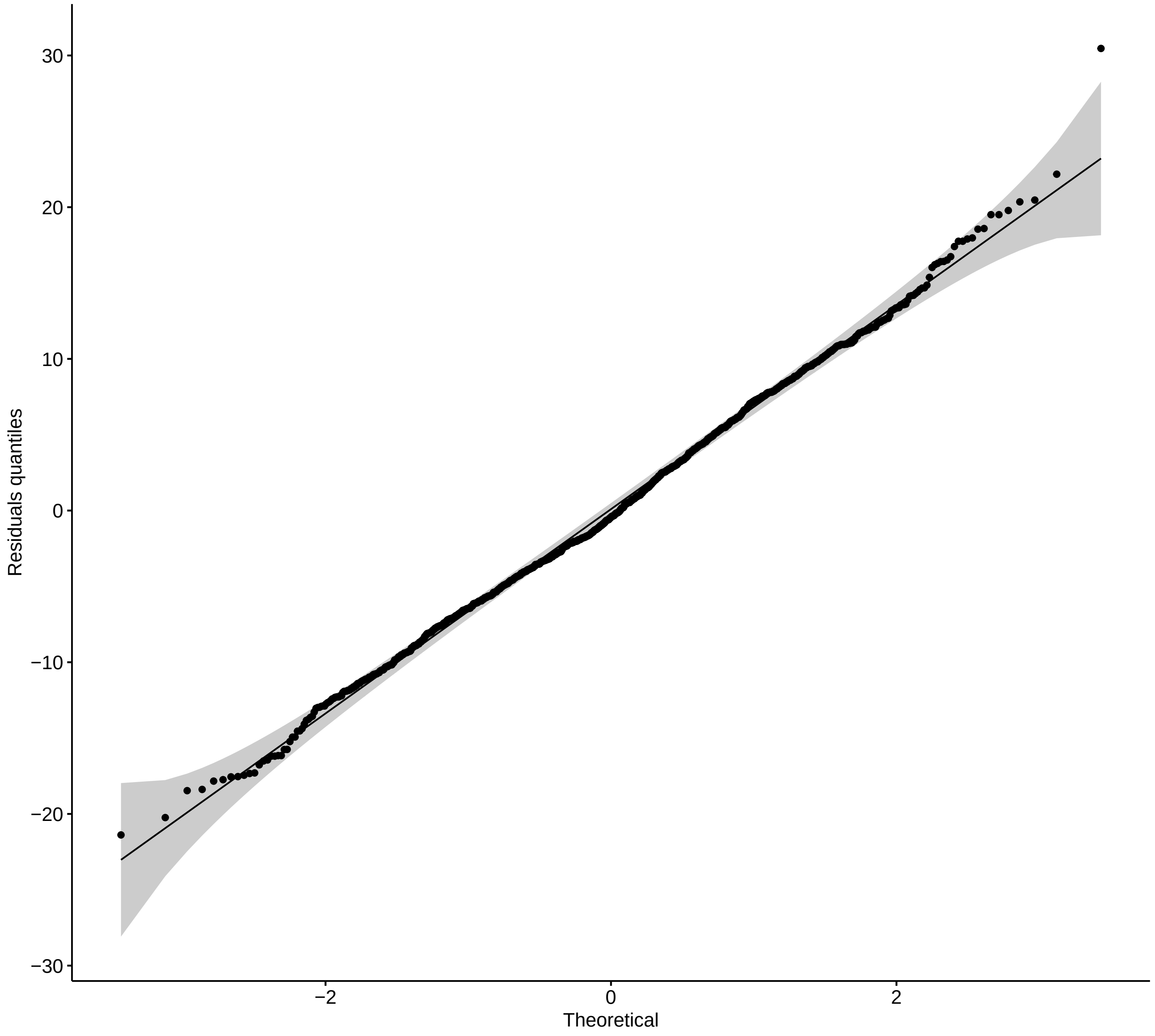


Supplementary Figure S2: Normal quantile plot versus GLS model residuals quantiles

##### 3.1.2 Standardised versus fitted plot

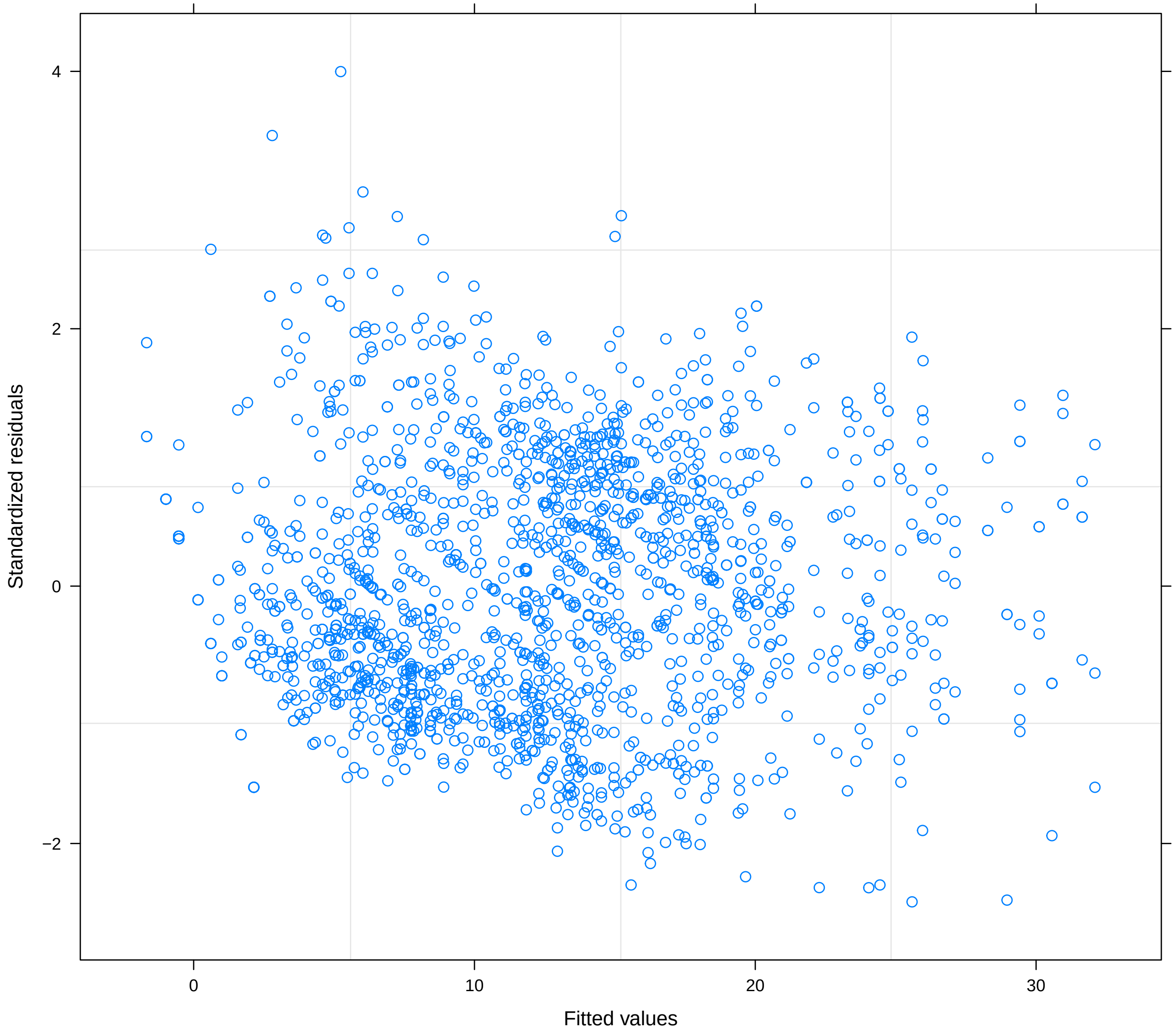


Supplementary Figure S3: Standardised residuals versus fitted values plot for the GLS model
